# A PTP1B-Cdk3 signaling axis promotes cell cycle progression of human glioblastoma cells through an Rb-E2F dependent pathway

**DOI:** 10.1101/2022.06.14.496178

**Authors:** Olga Villamar-Cruz, Marco Antonio Loza-Mejía, Alonso Vivar-Sierra, Héctor Iván Saldivar-Cerón, Genaro Patiño-López, Jonadab Efraín Olguín, Luis Ignacio Terrazas, Leonel Armas-López, Federico Ávila-Moreno, Sayanti Saha, Jonathan Chernoff, Ignacio Camacho-Arroyo, Luis Enrique Arias-Romero

**Affiliations:** Unidad de Investigación en Biomedicina (UBIMED), Facultad de Estudios Superiores-Iztacala, UNAM. Tlalnepantla, Estado de México 54090, Mexico; Unidad de Investigación en Reproducción Humana, Instituto Nacional de Perinatología-Facultad de Química, Universidad Nacional Autónoma de México (UNAM) 04510, Mexico City, Mexico; Design, Isolation, and Synthesis of Bioactive Molecules Research Group, Chemical Sciences School, Universidad La Salle-México, Benjamín Franklin 45, Mexico City 06140, Mexico; Carrera de Médico Cirujano, Facultad de Estudios Superiores Iztacala, UNAM, Tlalnepantla 54090, Mexico; Laboratorio de Investigación en Inmunología y Proteómica, Hospital Infantil de Mexico Federico Gómez, Mexico City 06720, Mexico; Laboratorio Nacional en Salud FES-Iztacala, Facultad de Estudios Superiores-Iztacala, UNAM. Tlalnepantla, Estado de México 54090, Mexico; Unidad de Investigación. Instituto Nacional de Enfermedades Respiratorias Ismael Cosío Villegas. Mexico City 14080, Mexico; Cancer Signaling and Epigenetics Program, Fox Chase Cancer Center. Philadelphia, PA 19111. USA

**Keywords:** Protein Tyrosine Phosphatase 1B, Cell Cycle, Cdk3, Cancer, Glioblastoma.

## Abstract

Protein tyrosine phosphatase 1B (PTP1B) plays a key role in developing different types of cancer. However, the molecular mechanism underlying this effect is unclear. To identify possible molecular targets of PTP1B that mediate its positive role in tumorigenesis, we undertook a SILAC-based phosphoproteomic approach, which allowed us to identify the Cyclin-dependent kinase 3 (Cdk3) as a novel PTP1B substrate. Molecular docking studies revealed stable interactions between the PTP1B catalytic domain and Cdk3. In addition, we observed that PTP1B dephosphorylates a Cdk3 derived peptide at Tyrosine residue 15 *in vitro* and interacts with endogenous Cdk3 in the nucleus and cytoplasm of human glioblastoma (GB) cells. Finally, we found that the pharmacological inhibition of PTP1B or its depletion with siRNA leads to cell cycle arrest with the diminished activity of Cdk3, the consequent hypophosphorylation of Rb, and the down-regulation of E2F and its target genes Cdk1, Cyclin A, and Cyclin E1. These data delineate a novel signaling pathway from PTP1B to Cdk3 required for efficient cell cycle progression in an Rb-E2F dependent manner in human GB cells and suggest new therapeutic strategies for treating these tumors.

## Introduction

Glioblastomas (GB) or grade IV gliomas are the most common and aggressive malignant primary tumors in the Central Nervous System (CNS), with an incidence rate of 3.23 per 100,000 persons and a median age of 64 years old [1]. GB patients have an overall survival rate of 15 months, even when receiving the standard therapy consisting of the maximum bearable surgical removal followed by radiotherapy and chemotherapy with temozolomide [2]. Therefore, it is necessary to understand better the molecular mechanisms underlying GB initiation and progression; and identify novel prognostic molecular markers to provide new therapeutic strategies. Recent studies on GB biology have revealed multiple alterations in pro-survival and anti-apoptotic signaling pathways. These include the Ras/RAF/MEK/ERK, PI3K/Akt/mTOR, and Src/Fak networks [3–6].

PTP1B is a classical non-transmembrane protein tyrosine phosphatase that plays a key role in metabolic signaling and is a promising drug target for type 2 diabetes and obesity [7–9]. Recent evidence indicates that PTP1B is overexpressed in different types of cancer. However, the role of PTP1B in GB development remains unclear [10]. For instance, it has been reported that PTP1B might be involved in the progression of different types of tumors by dephosphorylating and activating numerous oncogenic substrates. Moreover, several groups have recently shown that PTP1B dephosphorylates the inhibitory Y529 site in Src, thereby activating this kinase [11–13]. Another PTP1B substrate, p62^Dok^, might be involved in the positive effects of PTP1B on tumorigenesis. When p62^DOK^ is phosphorylated, it forms a complex with p120^RasGAP^ leading to decreased Ras/RAF/MEK/ERK activity [14,15]. In contrast, when PTP1B dephosphorylates p62^Dok^, it can no longer bind to p120^RasGAP^ with the consequent hyperactivation of Ras/RAF/MEK/ERK signaling, promoting cell proliferation and tumor growth. However, tissue samples from PTP1B-deficient breast cancer mouse models gave inconsistent results regarding the tyrosine phosphorylation status of p62^Dok^ [16,17]. For all the reasons above, it is essential to identify novel targets of PTP1B that could account for its pro-tumorigenic effects.

We identified novel molecular targets of PTP1B that could mediate its promoting role in cancer using a SILAC-based strategy. This proteomic approach allowed us to identify more than 200 potential novel targets of PTP1B. This work focused on the Cyclin-dependent kinase 3 (Cdk3), a protein involved in G0/G1 reentry and into G1/S transition in mammalian cells through the specific phosphorylation of Rb, which promotes its dissociation of the transcription factor E2F and activates the transcription of E2F responsive genes [18,19]. Different research groups have shown that in Cdks 1, 2, 3, and 5, Tyr15 phosphorylation abrogates its kinase activity [20–22]; and that CDC25 phosphatases activate Cdks through dephosphorylation of the Thr14 and Tyr15 residues in their ATP binding loop [23,24]. However, the identity of the CDC25 family member responsible for the dephosphorylation and activation of Cdk3 remains unknown. Our results from the phosphoproteomics experiment point to the possibility that Cdk3 might be activated by PTP1B in a CDC25-independent manner. To test this hypothesis, we first performed local docking and molecular dynamics studies that revealed stable interactions between the PTP1B catalytic domain and Cdk3. In addition, *in vitro* phosphatase assays confirmed that a phosphopeptide corresponding to the residues 9-21 of Cdk3 is dephosphorylated by PTP1B at Tyr15. Next, we observed that PTP1B interacts with Cdk3 in the nucleus and cytoplasm of human GB cell lines. Finally, we found that the pharmacological inhibition of PTP1B with the allosteric inhibitor claramine, a small-molecule inhibitor that targets the non-conserved C-terminal tail of PTP1B [25], or its depletion with siRNAs leads to cell cycle arrest with the diminished activity of Cdk3 and its downstream effectors Rb and E2F. These data delineate a novel signaling pathway from PTP1B to Cdk3 required for efficient cell cycle progression in an Rb-E2F dependent manner in human GB cells.

## Results

### Loss of PTP1B Leads to Increased Phosphorylation of Proteins Involved in Cell Proliferation

PTP1B occupies a central position in oncogenic signaling. In the last decade, several substrates of this phosphatase have been validated [26–28]. However, additional substrates that mediate the oncogenic effects of PTP1B in different types of cancer remain to be determined. In order to identify potential PTP1B substrates that might explain its promoting role in tumorigenesis, we used a quantitative proteomics approach based on high-resolution LC-MS upon differential labeling of wild-type and KO fibroblasts [29] with SILAC media (Figs. 1A and 1B). This phosphoproteomic approach allowed the identification of more than 200 potential direct or indirect targets of PTP1B with diverse cellular functions, including cell signaling, cell cycle progression and proliferation, metabolism, differentiation, apoptosis, and transcription (Fig. 1C and Supplemental Tables S1 and S2). Among the proteins predicted to be PTP1B enzymatic targets, we focused on those known to be involved in tumorigenesis (Table 1). In particular, we decided to investigate further Cdk3, a protein involved in G0/G1 reentry and into G1/S transition in mammalian cells that recently has been linked to the development of breast, colon, and esophageal cancer [19,30,31].

**Table 1.** Identification of cell cycle related substrates of PTP1B.

**Figure 6**
| Protein Name | pY Site | Sequence | PTM Score | KO/wt Ratio |
| --- | --- | --- | --- | --- |
| EGFR | Tyr1172 | LDNPDYQQDFF | 83.7 | 1.09 |
| EphA | Tyr772 | DPEATYTTSGGKI | 148.55 | 1.52 |
| p38 | Tyr182 | DEMTGYVATRL | 114.49 | 1.6 |
| CTNND | Tyr904 | LDNNYSTPNER | 127.75 | 2.02 |
| Cdk3 | Tyr15 | KIGEGTYGVVYKA | 83.47 | 1.98 |

**Figure 1.**
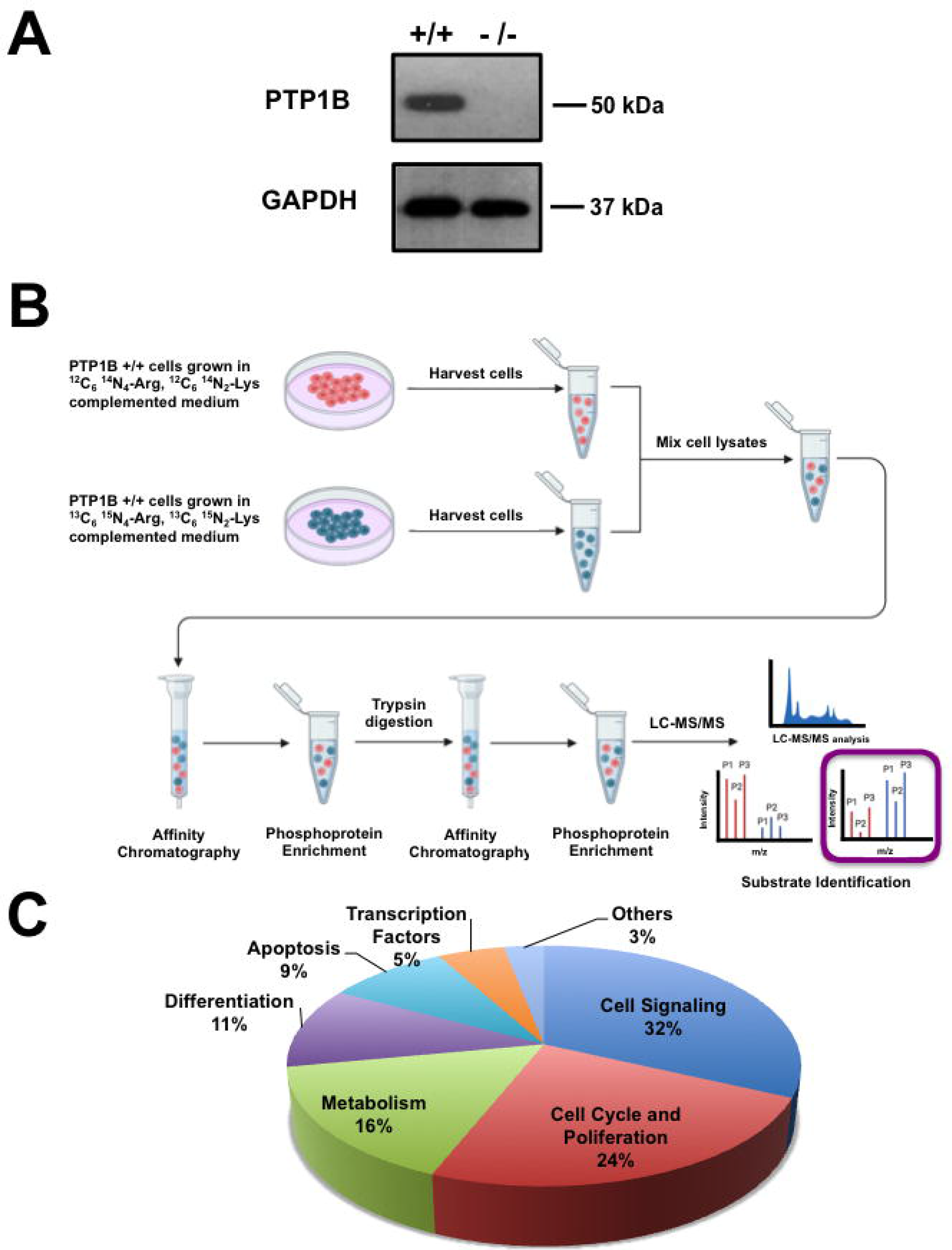
Schematic illustration of the SILAC-based quantitative phosphoproteomic approach. (A) Representative Western blot of PTP1B *+/+* and PTP1B −/− cells. (B) PTP1B *+/+* and PTP1B *−/−* cells were cultured in “light” (red) or “heavy” (blue) medium and stimulated with EGF for 15 min. After lysis, the samples were mixed and incubated with antiphosphotyrosine antibodies for enrichment of tyrosine-phosphorylated proteins. Then, the phosphoprotein fraction was digested with trypsin, and phosphopeptides were enriched again with TiO_2_ beads. Phosphopeptides were analyzed by LC-MS/MS. (C) Proteins identified in this assay were classified according to their cellular function.

### The Cyclin-dependent kinase 3 (Cdk3) Is a Novel Substrate of PTP1B

In order to determine if PTP1B dephosphorylates Cdk3 we performed peptide-protein docking experiments followed by molecular dynamics simulations. First, PEPstrMOD online platform [32] was used to predict the structure of the sequence of the Cdk3 peptide (KIGEGT<u>pY</u>GVVYKA), where the underlined residue corresponds to phosphoTyr15. Next, we generated the interaction model by docking in the ClusPro online server [33,34], using the crystal structure of PTP1B catalytic domain (PDB ID: 2HNP) [35]. The image of the predicted complex shows that the phosphorylated Tyr15 of Cdk3 is in close proximity to the catalytic residue Cys215 located in the phosphatase domain of PTP1B, with a predicted distance of 3.4 Å, being within the distance range for the transition state of the dephosphorylation reaction (Supplementary Figs. 1A and 1B). Motivated by these results, we decided to perform protein-protein molecular docking studies with the complete structures of PTP1B and Cdk3. Since Cdk3 crystallographic structure has not been reported, its full structure was generated by homology modeling using Cdk2, (PBD ID: 1B38) [36] as a template due to the high identity between these two enzymes. Our results indicated that the ligand-binding energy for the complex was −1723.932 kcal/mol, and the catalytic residues Cys215 and Arg221 on PTP1B’s phosphatase domain were in close proximity to the phosphorous atom of Cdk3 phospho-Tyr15, with an estimated minimal distance of 4.21 Å (Table 2 and Figs. 2A and 2B). In the proposed model, some basic residues such as Asp38 and Glu12 of Cdk3 interact with Arg 221, a critical residue of the catalytic site of PTP1B. This interaction seems to bring closer pTyr15 to the catalytic residue Cys 215 in PTP1B (Supplementary Fig. 1C). Similar values were obtained for a protein-protein docking model of PTP1B in complex with the IR β and DOK1, two well-characterized substrates of this phosphatase; and with the ephrin receptor EphA, one of the novel potential PTP1B substrates identified in our SILAC assay (ligand binding energies of −1677.228 kcal/mol, −1616.39 kcal/mol and −1304.71 kcal/mol respectively, minimal distance from the catalytic residues on PTP1B’s phosphatase domain to the phosphorous atom of IR β phospho-Tyr1146, 4.18 Å, to the phosphorous atom of DOK1 phospho-Tyr362, 4.18 Å, and to the phosphorous atom of EphA phospho-Tyr772, 4.45 Å respectively. Figs. 2C-2G and Table 2). These results suggest that Cdk3 should be a suitable substrate for PTP1B. In addition, the plot of the RMSD variations indicates that the complexes predicted by molecular docking have a stable conformation during the simulation. The analysis of the variations in the distance between PTP1B and Cdk3 showed that it is lower than 2.5 Å, suggesting that PTP1B forms a stable complex with Cdk3 and theoretically can dephosphorylate it (Supplementary Fig. 1D).

**Table 2.** Minimal distance from the catalytic residues on PTP1B’s phosphatase domain to the phosphorous atom of its substrates.

| Model | Initial distance<br>Cys215-pTyr (Å ) | Minimum distance<br>Cys215-pTyr after<br>MD simulation (Å ) | Ligand Binding<br>Energy<br>(kcal/mol) |
| --- | --- | --- | --- |
| PTP1B - Cdk3 | 6.93 | 4.21 | -1723.93 |
| PTP1B - IR $\beta$ | 5.41 | 4.18 | -1677.23 |
| PTP1B - DOK1 | 4.56 | 4.18 | -1616.39 |
| PTP1B - EphA | 4.46 | 4.45 | -1304.71 |

**Figure 2.**
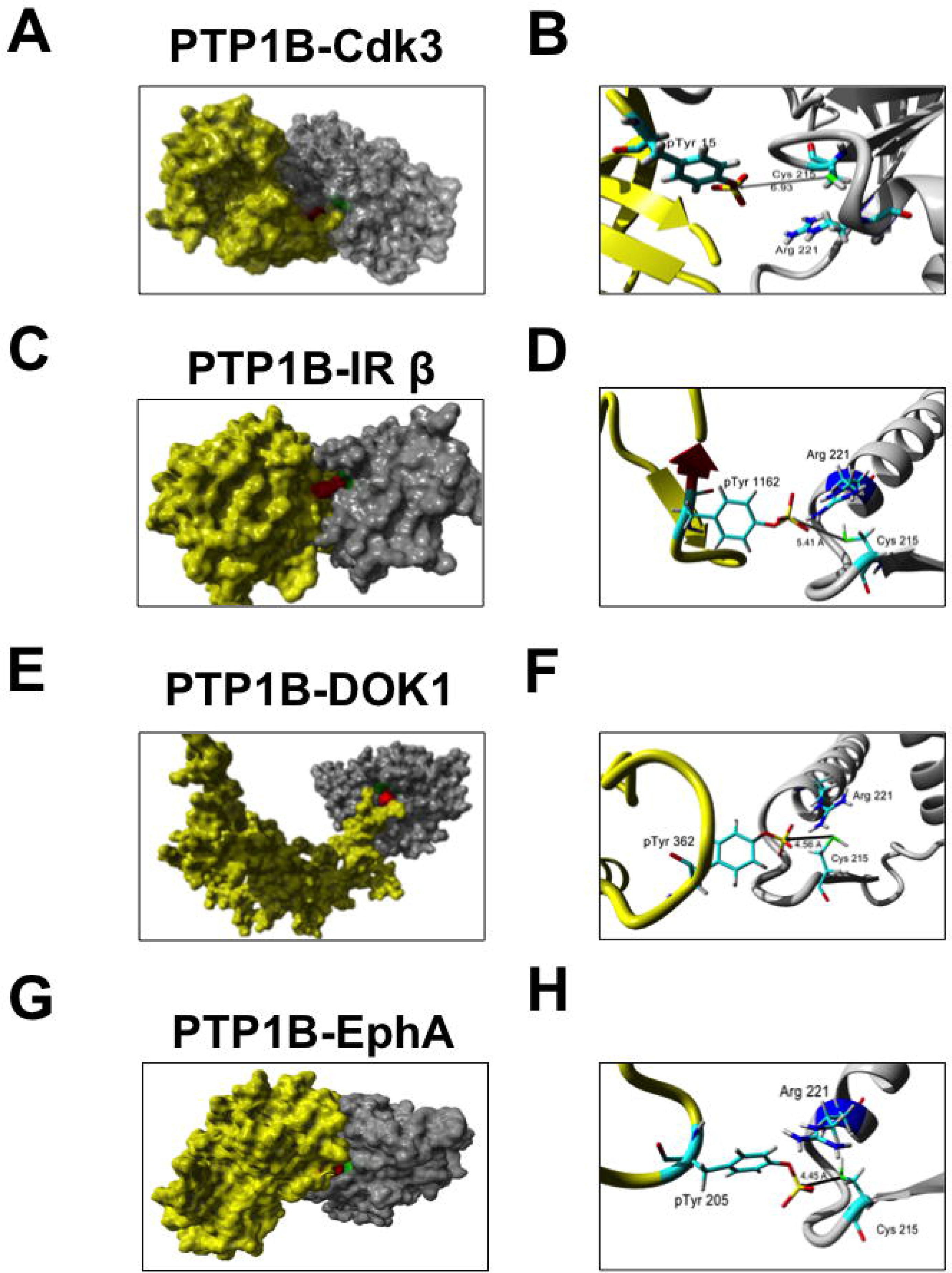
Predicted structure of PTP1B in complex with some of the putative substrates identified by SILAC-based phosphoproteomics. (A) Visualization of the complex of Cdk3 (yellow) and the catalytic domain of PTP1B (grey). The catalytic residue Cys215 of PTP1B and pTyr15 of Cdk are indicated in green and red, respectively. (B) Closer view of the PTP1B-Cdk3 interaction. The distance between pTyr15 of Cdk3 and Cys215 of PTP1B is indicated. (C) Visualization of the complex of IR β (yellow) and PTP1B (grey). The catalytic residue Cys215 indicated in green is close to IR β pTyr1146 indicated in red. (D) Closer view of the PTP1B-IR β interaction. The distance between Cdk3 pTyr1146 and PTP1B Cys215 is indicated. (E) Visualization of the complex of DOK1 (yellow) and PTP1B (grey). The catalytic residue Cys215 indicated in green is close to DOK1 pTyr362 indicated in red. (F) Closer view of the PTP1B-DOK1 interaction. The distance between DOK1 pTyr362 and PTP1B Cys215 is indicated..(G) Visualization of the complex of EphA (yellow) and PTP1B (grey). The catalytic residue Cys215 indicated in green is close to EphA pTyr205 indicated in red. (F) Closer view of the PTP1B-EphA interaction. The distance between EphA pTyr205 and PTP1B Cys215 is indicated.

Next, we explored if Cdk3 is a PTP1B substrate using an *in vitro* phosphatase assay. To this end, purified PTP1B was incubated with a phosphopeptide corresponding to residues 9-21 of Cdk3 or a phosphopeptide derived from the IR β as a positive control. We observed that PTP1B dephosphorylated both peptides with the same efficiency, suggesting that Cdk3 is a substrate of PTP1B (Fig. 3A).

**Figure 3.**
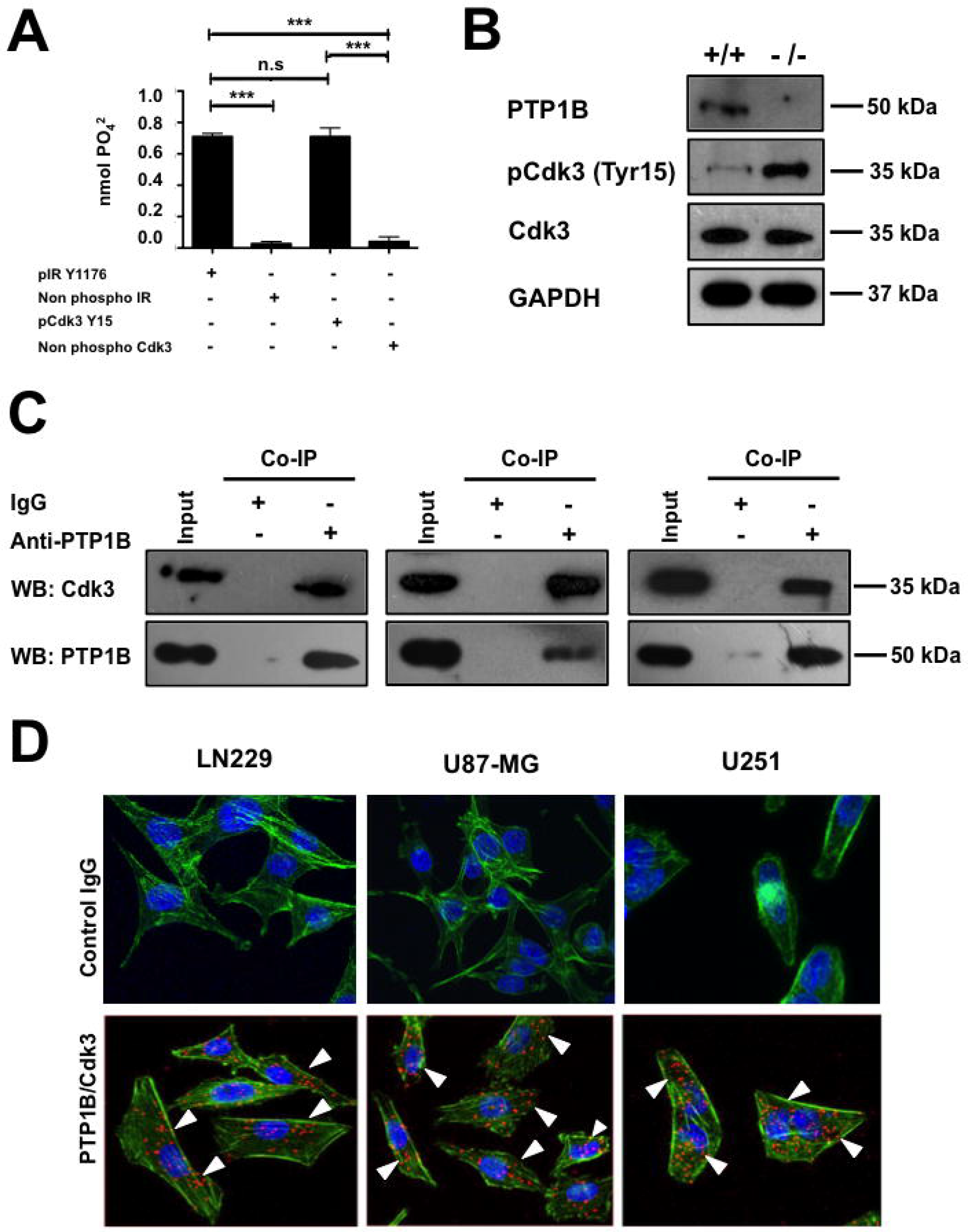
PTP1B dephosphorylates a Cdk3 derived peptide *in vitro*, and both proteins interact in GB cells. (A) The *in vitro* activity of PTP1B against a phosphopeptide corresponding to residues 9-21 of Cdk3 (KIGEGTpYGVVYKA) was determined by using a Malachite-Green based assay. An IR β phosphopeptide (TRDIpYETDYYRK) was used as positive control and the non-phosphorylated peptides of Cdk3 and IR β were used as negative controls. Each assay was independently repeated with four replicates and the nmol of phosphate released were calculated. Data are presented as means ± S.E. (B). Assessment of Cdk3 Tyr15 phosphorylation in PTP1B *+/+* and PTP1B −/− MEFs. Whole cell lysates were subjected to Western blot with anti-PTP1B or anti-Cdk3 antibodies. Total Cdk3 was immunoprecipitated and the detection of inactive Cdk3 was assessed by immunoblot using anti-phospho Cdk1/2/3 Tyr15 antibodies. GAPDH was used as loading control. (C) Co-immunoprecipitation of endogenous PTP1B and Cdk3. LN229, U87-MG and U251 cell lysates were subjected to immunoprecipitation with anti-PTP1B or isotype control IgG antibodies and the presence of Cdk3 in the complex was assessed with anti-Cdk3 antibodies. The presence of PTP1B and Cdk3 in cell extracts prior to immunoprecipitation was verified using specific antibodies (Input). (D) The physical interaction between endogenously expressed PTP1B and Cdk3 in LN229, U87-MG and U251 GB cells was determined using proximity ligation assays. The figure shows representative confocal microscopy images in which PTP1B-Cdk3 interactions appear as individual fluorescent red dots (lower panels). Anti-mouse and anti-rabbit isotype IgG antibodies were used as negative controls (upper panels). Scale bar, 50 μm.

### PTP1B Interacts with Cdk3 in a Cellular Context

Given that PTP1B dephosphorylates a Cdk3 derived peptide *in vitro*, we confirmed by immunoblot if PTP1B −/− fibroblasts display decreased Cdk3 activity. To this end, total Cdk3 was immunoprecipitated from PTP1B +/+ and PTP1B −/− MEFs and inactive Cdk3 was detected with anti-phospho Cdk1/2/3 (Tyr15) antibodies (Fig. 3B). Next, we tested if both proteins interact in a cellular context. To this end, we examined the physical association between endogenous PTP1B and Cdk3 in human GB cells by co-immunoprecipitation assays. LN229, U87-MG and U251 cells were lysed and PTP1B was immunoprecipitated with anti-PTP1B antibodies. The immunoprecipitates were separated by acrylamide gel electrophoresis and probed for associated Cdk3 by Western blot analysis. The presence of Cdk3 was readily detectable upon probing the immunoblots with the anti-Cdk3 antibody in immunoprecipitates (Fig. 3C). Next, we confirmed the interaction of endogenous PTP1B and Cdk3 in LN229, U87-MG, and U251 GB cells by proximity ligation assays using the corresponding two primary antibodies raised in different species. Then, the cells were incubated with species-specific secondary antibodies attached to a unique DNA strand (PLA probes). If the PLA probes are located less than 40 nm apart in the cell, the DNA strands can interact, forming a circle that can be amplified by DNA polymerase. Hybridization with complementary fluorescent oligonucleotide probes allows the visualization of PTP1B-Cdk3 interactions as fluorescent red dots. The results revealed multiple loci of interactions between endogenous PTP1B and Cdk3 in the cytoplasm and nucleus in LN229, U87-MG, and U251 cells. Mouse and rabbit IgGs were used as negative controls (Figure 3D).

Altogether, our results suggest that Cdk3 is a novel substrate of PTP1B and that both proteins physically interact in a cellular context.

### Inhibition of PTP1B Induces G1/S Arrest and Impairs Cdk3 Activity in Human Glioblastoma Cells

To further understand the biological relevance of PTP1B-Cdk3 interaction, we analyzed the effect of PTP1B pharmacological inhibition on Cdk3 Tyr15 phosphorylation levels and in cell cycle progression. Firstable, LN229, U-87 MG, and U-251 cells were synchronized by serum deprivation for 48 hours. Then, the cells were treated with vehicle or 2 μM of claramine during 3 hours and the cell cycle arrest was released by the addition of serum. The cells were lysed at the indicated time points, total Cdk3 was immunoprecipitated and inactive Cdk3 was detected with anti-phospho Cdk1/2/3 (Tyr15) antibodies. Our results indicated that 12 hours after serum stimulation Cdk3 is non-phosphorylated in vehicle treated cells, whereas it remains on its inactive phosphorylated form in claramine treated cells, suggesting that PTP1B inhibition impairs Cdk3 activation (Figs. 4A and 4B). Since Cdk3 is involved in G0/G1 reentry and into G1/S transition in mammalian cells [18,19], we hypothesized that PTP1B inhibition might lead to a decrease in Cdk3 activity with the consequent cell cycle arrest in these phases. To test this idea, LN229, U87-MG, and U251 cells were synchronized by serum deprivation for 48 hours, incubated with vehicle, 2 μM of claramine, or transfected with non-targeting siRNAs or with siRNAs targeting PTP1B or Cdk3 (Supplementary Fig. 2A), and collected every 12 hours after the release of cell cycle arrest. Our results indicated that human GB cells treated with claramine presented a marked delay in the transition to the S phase compared to vehicle-treated cells, whereas PTP1B and Cdk3 depleted cells showed an arrest in G0/G1 and reduced proliferation, suggesting that PTP1B and Cdk3 are essential regulators of cell cycle progression in human GB cells (Fig. 4C and Supplementary Fig. 2B).

**Figure 4.**
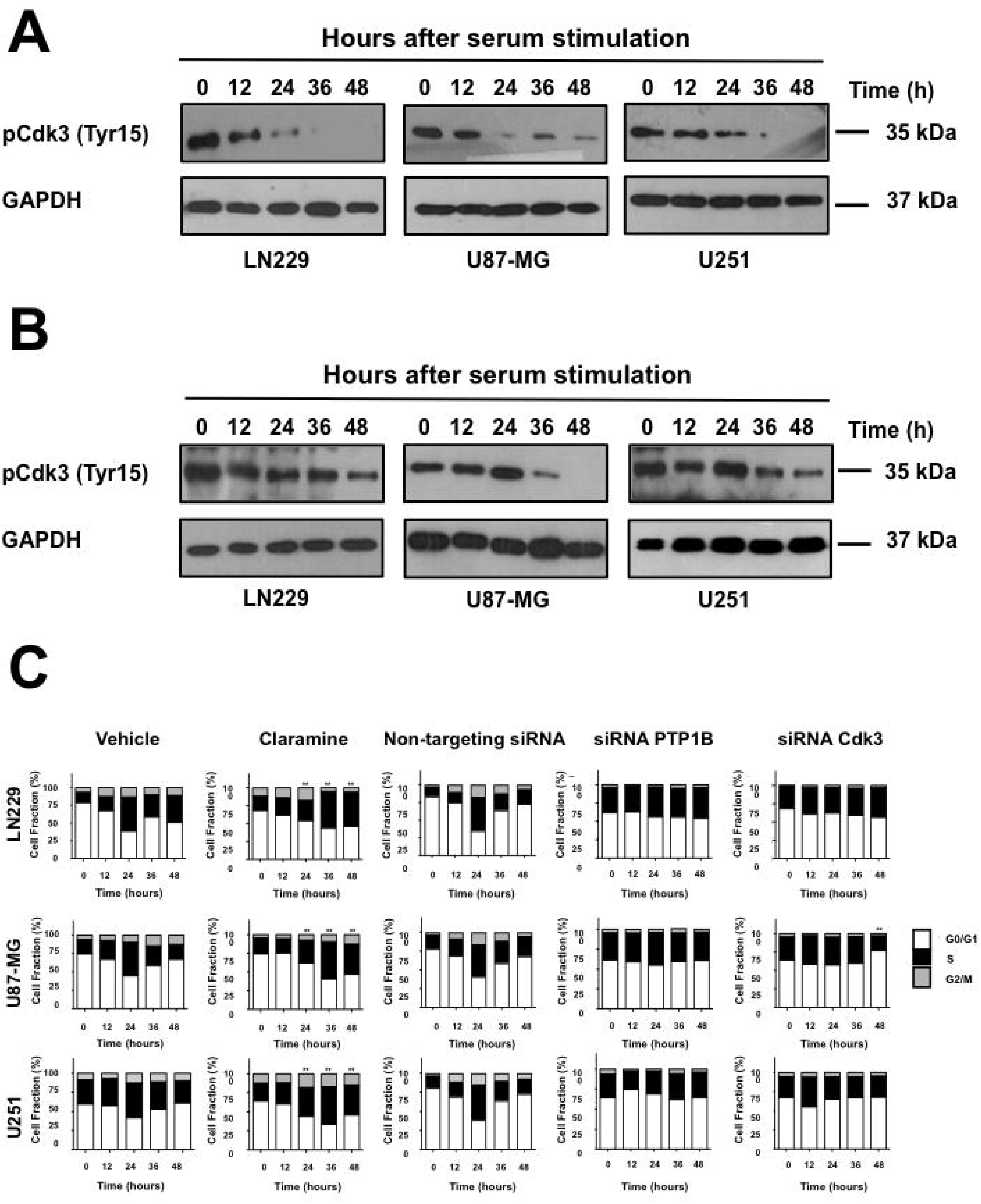
PTP1B reduced activity impairs Cdk3 dephosphorylation and delays cell cycle progression in human GB cells. GB cells were synchronized at G0 by serum deprivation for 48 hours and incubated 3 hours with vehicle (A) or claramine 2 μM (B). Cell cycle arrest was released by the addition of 10% FBS and cells were collected at the indicated times. Cdk3 was immunoprecipitated at the indicated times and the detection of inactive Cdk3 was assessed by immunoblot using anti-phospho Cdk1/2/3 Tyr15 antibodies. GAPDH was used as loading control using whole cell lysates before Cdk3 immunoprecipitation. (C) GB cells were synchronized at G0 and incubated 3 hours with vehicle, claramine 2 μM as previously mentioned, or transfected with siRNAs targeting PTP1B or Cdk3. Cell cycle arrest was released by the addition of 10% FBS, cells were fixed at indicated time points and stained with propidium iodide. Quantification of the percentage of cells at each phase is represented in the stacked graphics. Cells at G0/G1 phase are indicated in white boxes, cells at S phase in black boxes, and cells at G2/M in grey ones. Statistical differences in cells entering into S phase between control and claramine treated cells are indicated (*p<0.05) Data are representative of three independent experiments.

We next proved the effect of PTP1B inhibition on Cdk3 Tyr15 phosphorylation levels and in the activation of its downstream effectors Rb and E2F in human GB cells. LN229, U87-MG, and U251 cells were serum-starved during 48 hours, incubated for 3 hours in the presence of vehicle or claramine and stimulated with 10% of FBS. Cells were collected, Cdk3 was immunoprecipitated and the phosphorylation levels of Cdk3 Tyr15 were analyzed by Western blot using anti-phospho Cdk1/2/3 Tyr15 antibodies as previously described. Rb phosphorylation levels were assessed by Western blot using whole cell lysates. Our results indicate that PTP1B inhibition prevented the dephosphorylation of Cdk3 Tyr15 with the consequent hypophosphorylation of its downstream target Rb (Fig. 5A). Similarly, PTP1B knock down with siRNAs had a similar effect on Cdk3 and Rb phosphorylation levels (Fig. 5B). Considering that the phosphorylation of Rb is crucial to E2F transcription factor activity, we analyzed the expression levels of some E2F target genes necessary for G1/S transition by RT-qPCR. We observed that PTP1B inhibition repressed the expression of Cyclin A, Cyclin E1, and Cdk1 in all GB cell lines (Fig. 5C). Together, our results suggest that PTP1B regulates G1/S transition and promotes the expression of E2F downstream targets through the activation of Cdk3-Rb signaling, supporting the view that PTP1B is crucial for cell cycle regulation in GB (Fig. 6).

**Figure 5.**
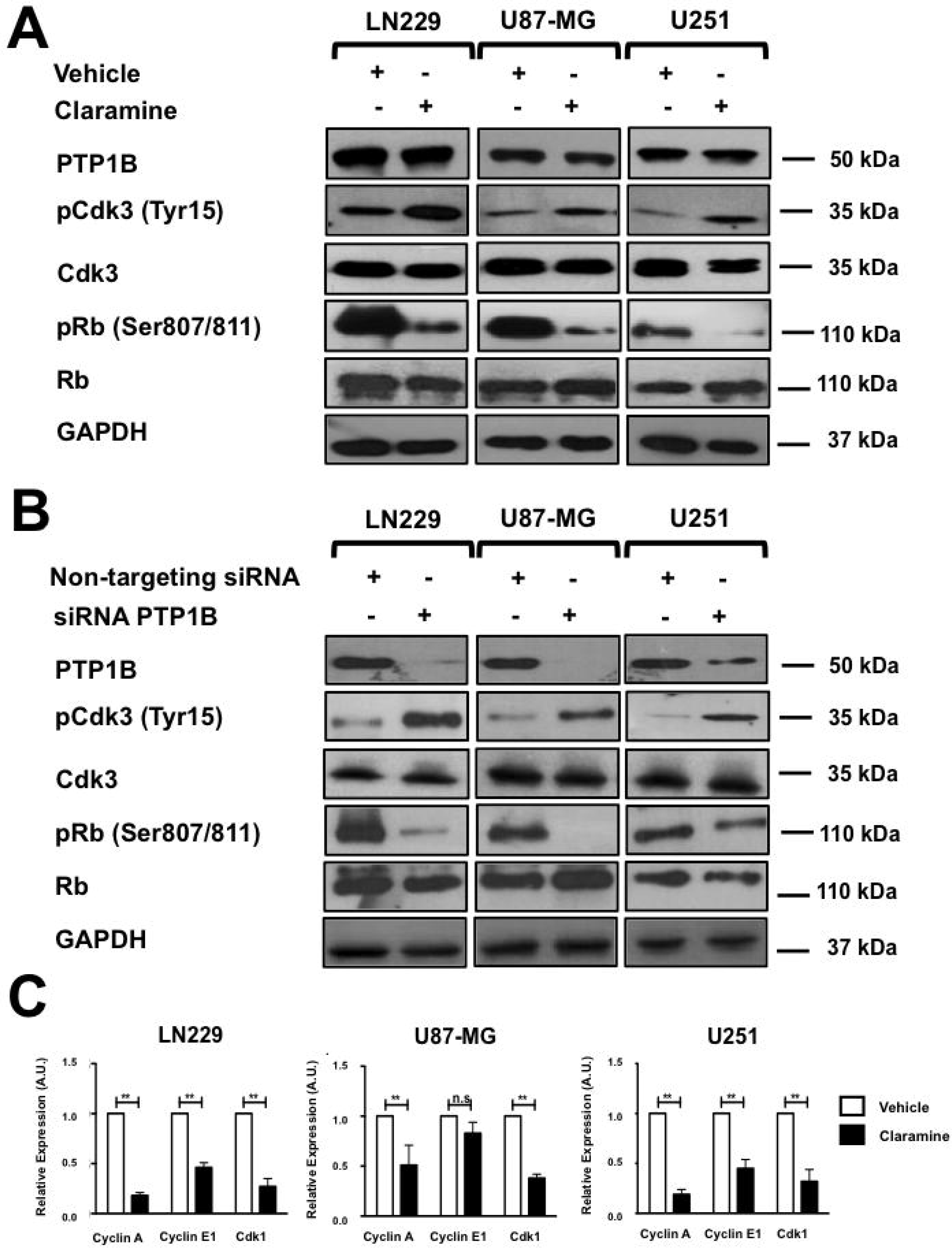
PTP1B reduced activity negatively affects Cdk3/Rb/E2F signaling in human GB cells. (A) The phosphorylation levels of Cdk3 and Rb in GB cells treated with vehicle or claramine 2 μM were assessed by immunoblot using total and phospho-specific antibodies. (B) The phosphorylation levels of Cdk3 and Rb in GB cells transfected with siRNAs targeting PTP1B were assessed by immunoblot using total and phospho-specific antibodies. (C) The expression levels of the E2F target genes: Cdk1, Cyclin A, and Cyclin E1 were assessed by RT-qPCR. All data were normalized to control GAPDH. Fold changes were calculated using the ΔCt method (2^−^ΔΔCt). Data are representative of three independent experiments.

**Figure 6.**
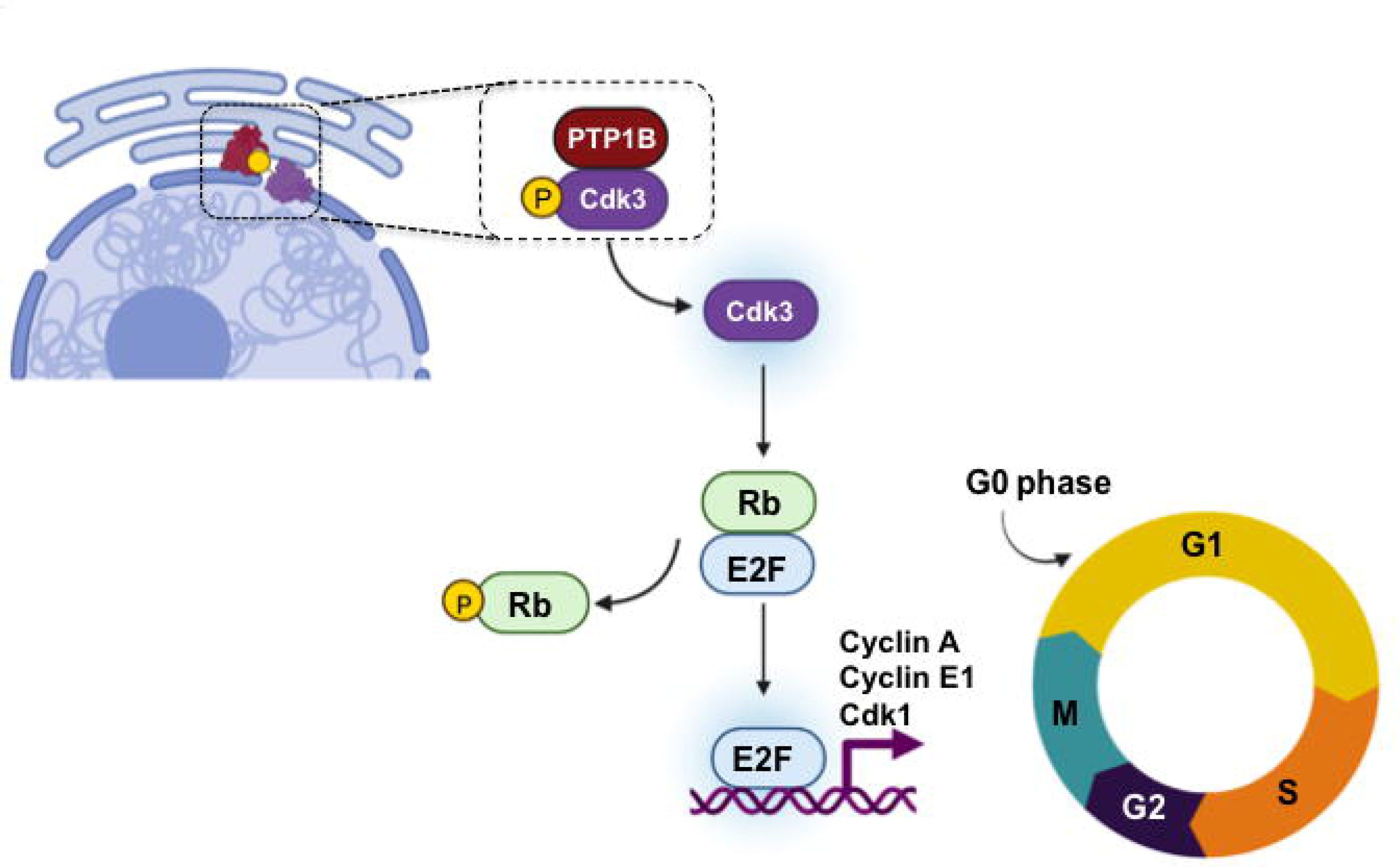
Proposed model for PTP1B regulation of cell cycle. PTP1B dephosphorylates and activates Cdk3, which in turn phosphoryates Rb, favoring its dissociation from E2F and promoting the expression of genes needed for the progression to S phase.

## Discussion

Recent evidence suggests that PTP plays an essential role in the development and progression of different types of cancer [10,37,38]. For example, *Ptpn1* gene is commonly amplified in breast, colon, prostate and gastric cancer [39–42]. In addition, recent reports indicate that PTP1B overexpression is associated with poor prognosis in several solid tumors, with a high PTP1B expression linked to poor outcome, including shortened progression-free and overall survival [42–45]. However, the molecular mechanisms that explain the contribution of PTP1B and its downstream targets to carcinogenesis are not entirely understood.

In this study, we used a SILAC-based phosphoproteomic approach to identify novel substrates of PTP1B that could explain its positive role in tumorigenesis. Our results indicate that Cdk3 is a new PTP1B target in GB cells. This conclusion is supported by our docking and molecular dynamics simulations, the ability of PTP1B to dephosphorylate *in vitro* a Cdk3 derived phosphopeptide, and to interact with it in the cytoplasm and nuclei of different human GB cell lines. Although, no differences were observed in the cellular processes evaluated in this work, the biological significance of a lower number of Cdk3-PTP1B interactions in U87-MG cells observed in the PLA assay needs to be evaluated. Furthermore, the pharmacological inhibition or knockdown with siRNAs of PTP1B promotes a significant delay in cell cycle progression *in vitro* and impairs Cdk3-Rb-E2F signaling. Therefore, PTP1B inhibition may represent an attractive therapeutic strategy for the treatment of GB. The effect of PTP1B inhibition observed in GB cells was not totally unexpected, since a previous report indicates that in pancreatic ductal adenocarcinoma cells, PTP1B knock down induces cell cycle arrest in G0/G1 phase with the consequent down-regulation of some cell cycle regulators needed for the G1/S transition such as Cdk2, Cdk4 and Cyclin D1 [46]. Furthermore, it has also been reported that PTP1B inhibition induces a G2 cell cycle arrest in renal cell carcinoma, pancreatic and hepatic cancer cells [47,48]. However, the molecular mechanism by which PTP1B regulates cell cycle progression in these cellular models has not been explored.

To our knowledge, this is the first report indicating that PTP1B regulates cell cycle progression through the activation of a Cdk. Finally, our results showed that PTP1B inhibition or depletion down-regulates the expression of some E2F genes required for G1/S transition, indicating that in our model, cell cycle progression may be regulated in part by a PTP1B/Cdk3/Rb/E2F signaling pathway. Our results suggest that inhibition of PTP1B should provide a new therapeutic strategy for the treatment of GB.

## Experimental procedures

### Antibodies and reagents

Antibodies used for western blot or PLA included anti PTP1B (sc-133258) from Santa Cruz Biotechnology (Dallas, TX. USA), Rb (#9309), phospho-Rb (Ser807/811) (#8516), and GAPDH (#5174) from Cell Signaling Technology (Boston, MA. USA). Anti-Cdk3 (ab191503) and anti phospho Cdk1/2/3 (Tyr15) (ab133463) from Abcam (Cambridge, MA. USA). Secondary antibodies conjugated to peroxidase (Goat anti-mouse cat #115-035-003 and Goat anti-rabbit cat #111-035-003) were purchased from Jackson ImmunoResearch Laboratory (West Grove, PA. USA). The allosteric PTP1B inhibitor claramine was purchased from Sigma-Aldrich (Burlington, MA, USA).

### Cell Culture

*Ptp1b*^−/−^ and *Ptp1b*^+/+^ MEFs were a gift from Dr. Benjamin Neel (New York University) [29]; these cells were maintained in DMEM (Gibco BRL, Walthman, MA, USA) supplemented with 10% FBS, 50 U/mL penicillin, and 50 μg/mL streptomycin. HEK293T, LN-229, U251, and U-87 MG cells were obtained from American Type Culture Collection (Manassas, VA, USA), and maintained in DMEM (Gibco BRL, Walthman, MA, USA) supplemented with 10% FBS, 50 U/mL penicillin, and 50 μg/mL streptomycin.

For SILAC experiments, *Ptp1b*^−/−^ and *Ptp1b*^+/+^ cells were grown in media containing ^12^C_6_,^14^N_4_-Arg and ^12^C_6_,^14^N_2_-Lys, or ^13^C_6_,^15^N_4_-Arg and ^13^C_6_,^15^N_2_-Lys (Thermo Fisher Scientific, Waltham, MA. USA) until the populations went through six passages. Then, cells were starved for 3 h prior to ligand stimulation with EGF (50 ng/mL, 10 min; PeproTech, Rocky Hill, NJ, USA), and were lysed in ice-cold lysis buffer (50 mM Tris-HCl, pH 7.5, 150 mM NaCl, 1% Nonidet P-40, 0.1% sodium deoxycholate, 1 mM EDTA, 1 mM sodium orthovanadate, 1 mM PMSF, 0.1 μg/mL aprotinin, 10 mM NaF) for 20 min. Lysates were precleared by centrifugation at 16,500 × *g* for 15 min. Protein amount determination was performed using the Bradford assay (Bio-Rad, Hercules, CA, USA). In SILAC experiments, cell lysates were mixed 1:1 (double labeling).

### Anti-phospho-Tyrosine Immunoprecipitation

For immunoprecipitation, 200 μg of anti-phospho-tyrosine 4G10 antibody (cat # ZMS16282. Sigma-Aldrich. St. Louis, MO. USA) were added together with 40 μL of protein A-Sepharose (GE Healthcare, Chicago, IL, USA) to mixed cell lysates containing up to 20 mg of total labeled proteins and incubated at 4 °C for 4 h. Precipitates were washed four times with lysis buffer, and precipitated proteins were eluted twice with urea buffer (7 M urea, 2 M thiourea, 50 mM HEPES, pH 7.5, 1% *n*-octyl glucoside) at 37 °C for 10 min.

### In-solution Protein Digestion

After anti-phospho-tyrosine immunoprecipitation, precipitated proteins were denatured in the urea buffer described above, and protein amount was measured using the Bradford assay (Bio-Rad, Hercules, CA, USA). Proteins were reduced by adding 2 mM DTT (final concentration) at 25 °C for 45 min, and thiols were carboxymethylated with 5.5 mM iodoacetamide at room temperature for 30 min. Endoproteinase Lys-C was added in an enzyme/substrate ratio of 1:100, and the proteins were digested at room temperature for 4 h. The resulting peptide mixtures were diluted with water to reach a final urea concentration below 2 M. For double digestion, modified trypsin (sequencing grade, Promega, Madison, WI, USA) was N added in an enzyme/substrate ratio of 1:100, and the digest was incubated at room temperature overnight. Trypsin activity was quenched by adding TFA to a final concentration of 1%.

### Titansphere Enrichment of Phosphopeptides

After trypsin digest, phosphopeptides were enriched using Titansphere chromatography (TiO_2_) columns as previously described [49,50]. Peptide samples were diluted 1:6 with 30 g/L 2,5-dihydroxybenzoic acid (DHB) in 80% ACN, 0.1% TFA. 5 μg of TiO_2_ beads (GL Sciences Inc., Torrance, California, USA) were washed once with elution buffer (NH_3_ water in 20% ACN, pH 10.5) and equilibrated with washing buffer (50% ACN, 0.1% TFA). TiO_2_ beads were loaded with DHB by washing with loading buffer (6 g/L DHB in 15% ACN). Peptide samples were loaded onto TiO_2_ beads for 30 min at room temperature on a rotating wheel. Subsequently, beads were washed three times with washing buffer, and bound phosphopeptides were eluted twice with 50 μL of elution buffer at room temperature for 10 min. The eluates were filtered through STAGE Tips in 200-μL pipette tips. 30 μL of 80% ACN, 0.5% acetic acid was applied to the STAGE Tips after filtering [51], and the flow-through was combined with the filtered sample. The pH value of the sample was adjusted with TFA to a value of approximately pH 7, and the eluates were concentrated in a vacuum concentrator. Before MS analysis, 5% ACN and 0.1% TFA (final concentrations) were added to the samples.

### LC-MS/MS Analysis

Mass spectrometric analysis was performed by nanoscale LC–MS/MS using an LTQ-Orbitrap Velos instrument (Thermo Fisher Scientific, Waltham, MA. USA) coupled to an Ultimate U3000 (Dionex Corporation, Sunnyvale, CA, USA) via a Proxeon nanoelectrospray source (Proxeon Biosystems, Denmark). Peptides were separated on a 75 μM PepMap C18 nano column.

Data were acquired in data-dependent mode: full scan spectra were acquired with a resolution of 60◻000 after accumulation of 1◻000◻000 ions. The “lock mass” option was used to improve the mass accuracy of precursor ions. The ten most intense ions were fragmented by collision-induced dissociation (CID) with a normalized collision energy of 35% and recorded in the linear ion trap (target value of 5000) based on the survey scan and in parallel to the orbitrap detection of MS spectra. Peptide identification was performed using the MASCOT search engine.

### Model of the Binding Complex of PTP1B catalytic domain and Cdk3

To generate a model that could show the interaction of Cdk3 and its derived peptide (sequence: KIGEGT<u>pY</u>GVVYKA where the underlined pY corresponds to phospho-Tyr15), we decided to perform peptide-protein and protein-protein docking experiments followed by molecular dynamics simulations. First, PEPstrMOD online platform [32] was used to predict the structure of the sequence of Cdk3 phosphopeptide while Cdk3 full structure was generated using Cdk2 (PBD ID: 1B38) [36] as a template with the homology modeling module implemented as part of Yasara Structure v.18.4.24. The catalytic domain of PTP1B was also downloaded from the PDB with the accession code (PDB ID: 2HNP) [35].

ClusPro online server https://cluspro.bu.edu/ **[52–55]**was used for the docking experiments. ClusPro 2.0 generates conformations based on different desolvation and electrostatic potential; those conformations are categorized through clustering. Molecular dynamics simulations were performed in duplicate using Yasara Structure v. 18.4.24 [56,57] using the AMBER 14 force field using a previously reported protocol [58]. Briefly, each complex was embedded within a TIP3 water box with 10 Å to the box boundary. Periodic boundary conditions were considered. The temperature was set to 298 K, pH to 7.4, with the addition of sodium and chlorine ions for charge neutralization. A Particle Mesh Ewald (PME) algorithm with a cut-off radius of 8 Å was applied. Steepest descent energy minimization was performed, and then a total simulation time of 100 ns with a time step of 2.5 fs was carried out, recording snapshots at intervals of 500 ps. The analysis of the resulting trajectories was performed with a script included as part of Yasara software and included the root mean square deviation (RMSD) and the distance of the catalytic Cys215 and Arg221 on PTP1B’s phosphatase domain and the phosphorus atom on Cdk3 phospho-Tyr15.

For the models of IR β, DOK1 and EphA, the initial structures were retrieved from AlphaFold [59]. These structures were minimized using the same protocol in molecular dynamics as described before, for a total simulation time of 100 ns. Then, the complexes of PTP1B and these optimized structures were predicted using ClusPro under the same conditions as explained for the complexes with Cdk3.

### In vitro phosphatase assay

According to the manufacturer’s suggested protocol, the *in vitro* phosphatase assays were performed using the Protein Tyrosine Phosphatase 1B Assay Kit, Colorimetric (Sigma-Aldrich. St. Louis, MO. USA). We designed a phosphopeptide for the PTP1B-dependent phosphosite on Cdk3 (pY15: KIGEGT<u>pY</u>GVVYKA). In addition, we included a known PTP1B substrate peptide derived from the IR β (pY1146: TRDI<u>pY</u>ETDYYRK), as a positive control, and the non-phosphorylated peptides of Cdk3 and IR β as negative controls. These peptides were synthesized (95% purity, PROBIOTEK, San Nicolás de los Garza, NL, Mexico) and, prior to assays, and freshly diluted to 150◻μM final concentration in 1x Assay Buffer (150 mM NaCl, 50 mM MES, 1 mM DTT, 1 mM EDTA, 0.05% NP-40, pH 7.2). The reactions were performed in triplicate in a 96-well plate at 30°C, adding 75 ◻μM of each phosphopeptide and 2.5 ng of recombinant PTP1B for 30 min. The addition of Malachite green solution terminated the reactions, and the absorbance was measured at 620 nm on an Epoch 2 microplate reader (Tekan, Winoosky, VT, USA).

### Immunoblotting and Co-Immunoprecipitation

GB cells were lysed in RIPA buffer (20nM Tris-HCL pH 7.4, 150 nM NaCl, 1 mM EDTA, 1% Triton X-100, 0.5% SDS, 1% sodium deoxycholate, 1X protease inhibitor Cocktail and 1X PhosSTOP (Sigma-Aldrich. St. Louis, MO. USA). Immunoblots on Immobilon-P membranes (Millipore. Burlington, Massachusetts. USA) were blocked in 5% nonfat dried milk in TBS-Tween-20 0.5% or 1% BSA, incubated primary and secondary antibodies, and visualized using enhanced chemiluminescence reagents (ECL, Amersham Pharmacia. Buckinghamshire. UK). For co-immuniprecipitation cells were lysed for 30 min in PBSCM buffer (100 mM Na_2_PO_4_, 150 mM NaCl, pH 7.2, 1 mM CaCl_2_, 1 mM MgCl_2_ and 5 μM ATP), homogenized and centrifuged at 13,500rpm for 10 min at 4°C. The supernatants were recovered, clarified and incubated with primary antibodies or mouse or rabbit IgG isotype control antibodies for 4 h at 4°C, and incubated with Protein G Sepharose beads (GE Healthcare. Braunschweig, Germany). The immune complexes were washed three times with PBSCM buffer and the bound material was eluted using sample buffer for 5 min at 90 °C. The eluate was resolved on 10% SDS–PAGE and analyzed by immunoblot. All the antibodies were used at concentrations as recommended by the supplier.

### Proximity Ligation Assay

According to the manufacturer’s instructions, PLA experiments were performed using Duolink^®^ kit (DUO92101, Sigma-Aldrich, St. Louis, MO, USA.). A total of 4000 cells of each cell line were plated in 16 well chambers (178599, Nunc Lab-Teck, Thermo Scientific, Waltham, MA, USA) in DMEM medium with 10% fetal bovine serum and incubated at 37 °C under a 95% air and 5% CO_2_ atmosphere for 24 h. Cells were fixed with PBS/PFA 4% for 15 min and permeabilized with 0.5% Triton X−100 for 30 min. Then, cells were blocked with 40 μL of blocking solution in a humidity chamber at 37 °C for 1 h, and incubated with the primary antibodies: polyclonal rabbit antibody against PTP1B (2 μg/mL; #5311; Cell Signaling Technology, Boston, MA, USA) and monoclonal mouse antibody against Cdk3 (2 μg/mL; sc-81836; Santa Cruz Biotechnology, Dallas, TX, USA) at 4 °C overnight. To detect the primary antibodies, secondary proximity probes binding rabbit and mouse immunoglobulin (PLA probe rabbit PLUS and PLA probe mouse MINUS, Olink Bioscience, Sigma-Aldrich, St. Louis, MO, USA) were diluted 1:15 and 1:5 in blocking solution, respectively. Cells were then incubated with the proximity probe solution at 37 °C for 1 h, washed three times in 50 mM Tris pH 7.6, 150 mM NaCl, 0.05% Tween-20 (TBS-T), and incubated with the hybridization solution containing connector oligonucleotides (Olink Bioscience, Sigma-Aldrich, St. Louis, MO, USA) at 37 °C for 45 min. Samples were washed with TBS-T and subsequently incubated in the ligation solution at 37 °C for 45 min. The ligation solution contained T4 DNA ligase (Fermentas, Sigma-Aldrich, St. Louis, MO, USA), allowing the ligation of secondary proximity probes and connector oligonucleotides to form a circular DNA strand. Subsequently, the samples were washed in TBS-T and incubated with the amplification solution containing phi29 DNA polymerase (Fermentas, Sigma-Aldrich, St. Louis, MO, USA) for rolling circle amplification at 37 °C for 90 min, and washed three times with TBS-T. Finally, the samples were incubated with the detection mix solution containing Texas Red-labeled detection probes that recognize the amplified product (Olink Bioscience, Sigma-Aldrich, St. Louis, MO, USA) at 37 °C for 1 h, washed twice in SSC-T buffer (150 mM NaCl, 15 mM sodium citrate, 0.05% Tween-20, pH 7), and were coverslipped with a fluorescence mounting medium (Biocare Medical, Pacheco, CA, USA). Fluorescent signals were detected by laser scanning microscopy (Leica TCS SP8, Wetzlar, Germany), and PLA-positive signals were quantified using MetaMorph software.

### Cell Cycle Analyses

LN229, U87-MG and U251 cells (2 × 10^5^) were seeded in six-well cell culture plates, synchronized by serum deprivation for 48 h, incubated with the vehicle of 2 μM of claramine as previously reported [60], and collected at the indicated time points after release of cell cycle arrest. Cells were washed with PBS and fixed in 70% ethanol overnight. Cells were then washed twice with PBS and stained using a Propidium Iodide/RNase Staining kit (BD PharMingen. Franklin Lakes, NJ. USA) in the dark at room temperature for 30 min. Cell cycle analysis was carried out using an Attune NxT flow cytometer (Thermo Fisher Scientific, Waltham, MA. USA) and FlowJo software version 10.6. A total of 10,000 cells were collected for each sample for analysis.

### RT-qPCR

Total RNA was extracted from cells treated with claramine 5 μM or vehicle using RNeasy Mini kits, quantified by Nanodrop ND-1000, and reverse transcribed using the High Capacity cDNA Reverse Transcription Kit (Applied Biosystems). 1 ng cDNA was amplified by real-time PCR using Universal ProbeLibrary (UPL) probes (Roche). The primer sequences for real-time qPCR were: Cdk1 Fwd: 5’ ggaaaccaggaagcctagcatc 3’, Rev: 5’ ggatgattcagtgccattttgcc 3’; Cyclin A Fwd; 5’ tgtgcacggaaggactttgt 3’, Rev: 5’ tcctcttggagaagatcagccg 3’; Cyclin E1 Fwd: 5’ tgtgtcctggatgttgactgcc 3’, Rev: 5’ ctctatgtcgcaccactgatacc 3’; and GAPDH Fwd: 5’ ccccggtttctataaattgagc 3’, Rev: 5’ caccttccccatggtgtct 3’. UPL probes used were #80 for Cdk1, #69 for Cyclin A, #19 for Cyclin E1 and #63 for GAPDH. Each sample was run in 20 μL reaction using 2X FastStart Universal Probe Master with ROX (Roche). Reactions were performed in an ABI real-time PCR 7500 (Applied Biosystems, Foster City, CA). Ratios of mRNA levels to control values were calculated using the ΔCt method (2-ΔΔCt) at a threshold of 0.02 [61]. All data were normalized to control GAPDH. PCR conditions used: hold at 95°C for 10 min, followed by 40 cycles of 15 s at 95°C and 60 s at 60°C.

### Statistics

Statistical analysis was conducted using a two-way ANOVA or the unpaired Student *t*-test using the Prism software package (GraphPad Software. San Diego, CA, USA). Values of *p*<0.05 were considered significant.

## Acknowledgments

We thank Biol. Luis Enrique Florencio-Martinez for technical assistance and Dr. Norma Laura Delgado-Buenrostro for assistance in PLA confocal microscopy analysis. Olga Villamar-Cruz is a graduate student of Programa de Doctorado en Ciencias Bioquímicas, UNAM.

## Data Availability Statement

The original contributions presented in the study are included in the article; further inquiries can be directed to the corresponding author/s.

