## Supplemental Figure Legends for "A PTP1B-Cdk3 signaling axis promotes cell cycle progression of human glioblastoma cells through an Rb-E2F dependent pathway"

**Supplementary Figure 1. PTP1B-Cdk3 derived peptide Interaction Model.** (A) Visualization of the complex of Cdk3 phosphopeptide (grey) and PTP1B (yellow). The catalytic residue Cys215 indicated in blue is close to Cdk3 pTyr15. (B) Closer view of the PTP1B-Cdk3 interaction. The distance between Cdk3 pTyr15 and PTP1B Cys215 is indicated. (C) Closer view of the PTP1B-Cdk3 interaction. The proximity between pTyr15, Glu12 and Asp38 of Cdk3 to Cys215, Arg45 and Arg221 of PTP1B is showed. (D) The Root-Mean-Square Deviation (RMSD) for the variations in the distance of the sulfur atom of the Cys215 residue towards the phosphate group of phosphotyrosine residues $S_{(Cys)}$-$P_{(pTyr)}$ is shown. The plot of the RMSD variations shows that the complexes predicted by molecular docking reach convergence after a small equilibration period. The analysis of the variations in the $S_{(Cys)}$-$P_{(pTyr)}$ distance for the complex of phosphorylated Cdk3 and PTP1B show that distance could be close to 4 Å, suggesting that conformational changes allow the dephosphorylation of pTyr 15 in Cdk3 are feasible.

**Supplementary Figure 2. PTP1B and Cdk3 depletion impairs cell proliferation. (A)** GB cells were transfected with non-targeting siRNAs, siRNAs targeting PTP1B or Cdk3, and the expression of both proteins was assessed by immunoblot. GAPDH was used as loading control. (B) GB cells were synchronized at G0 by serum deprivation and incubated 3 hours with vehicle, claramine 2 μM or transfected with non-targeting siRNAs, siRNAs targeting PTP1B or Cdk3. Cells were counted every 24 hours. Proliferation of LN229, U87-MG and U251 cells is represented as mean ± S.E. of three independent experiments.
