## Supplementary figures and images for "A PTP1B-Cdk3 signaling axis promotes cell cycle progression of human glioblastoma cells through an Rb-E2F dependent pathway"

### Supplemental Figure 1

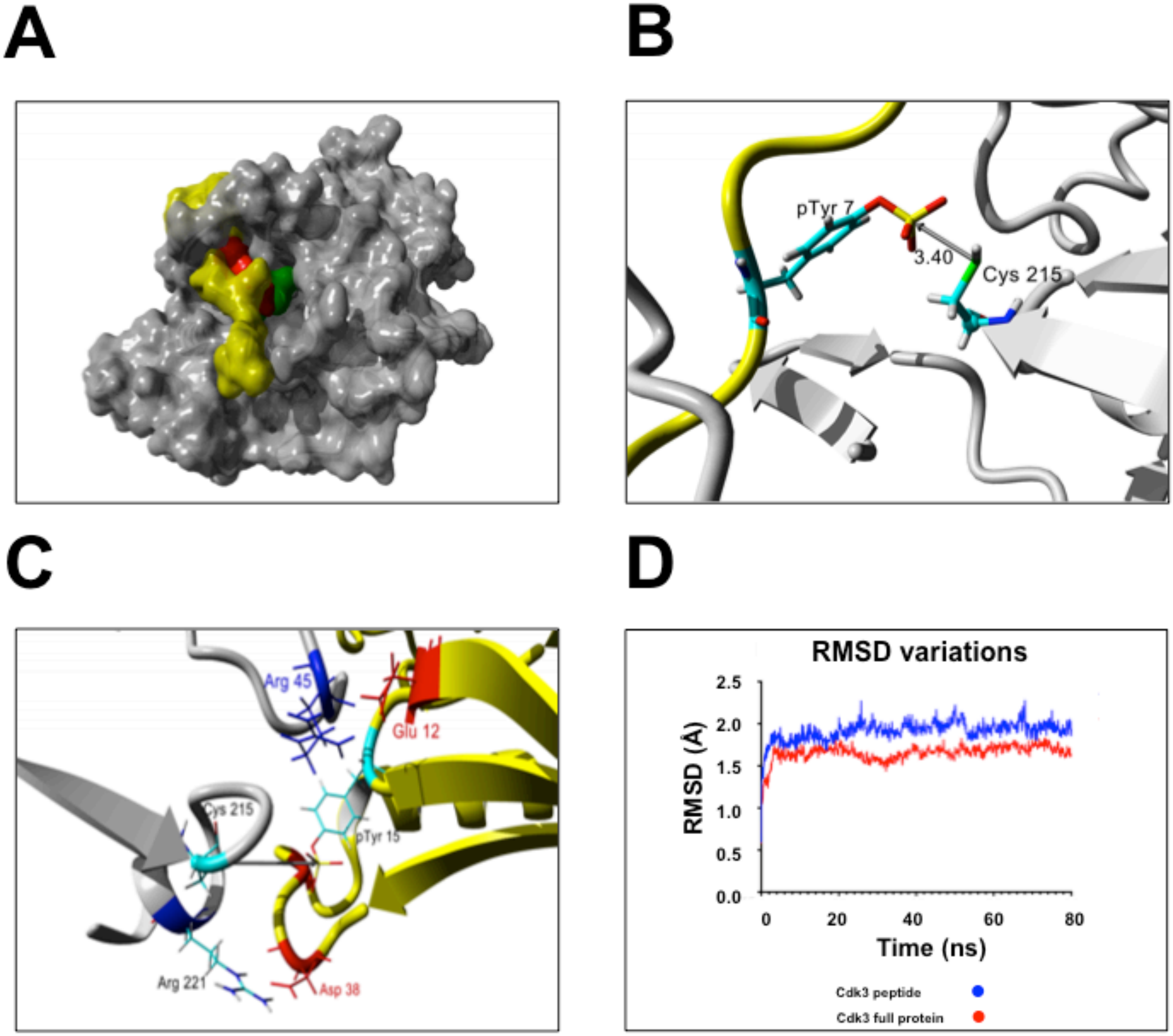


**Supplementary Figure 1**

### Supplemental Figure 2

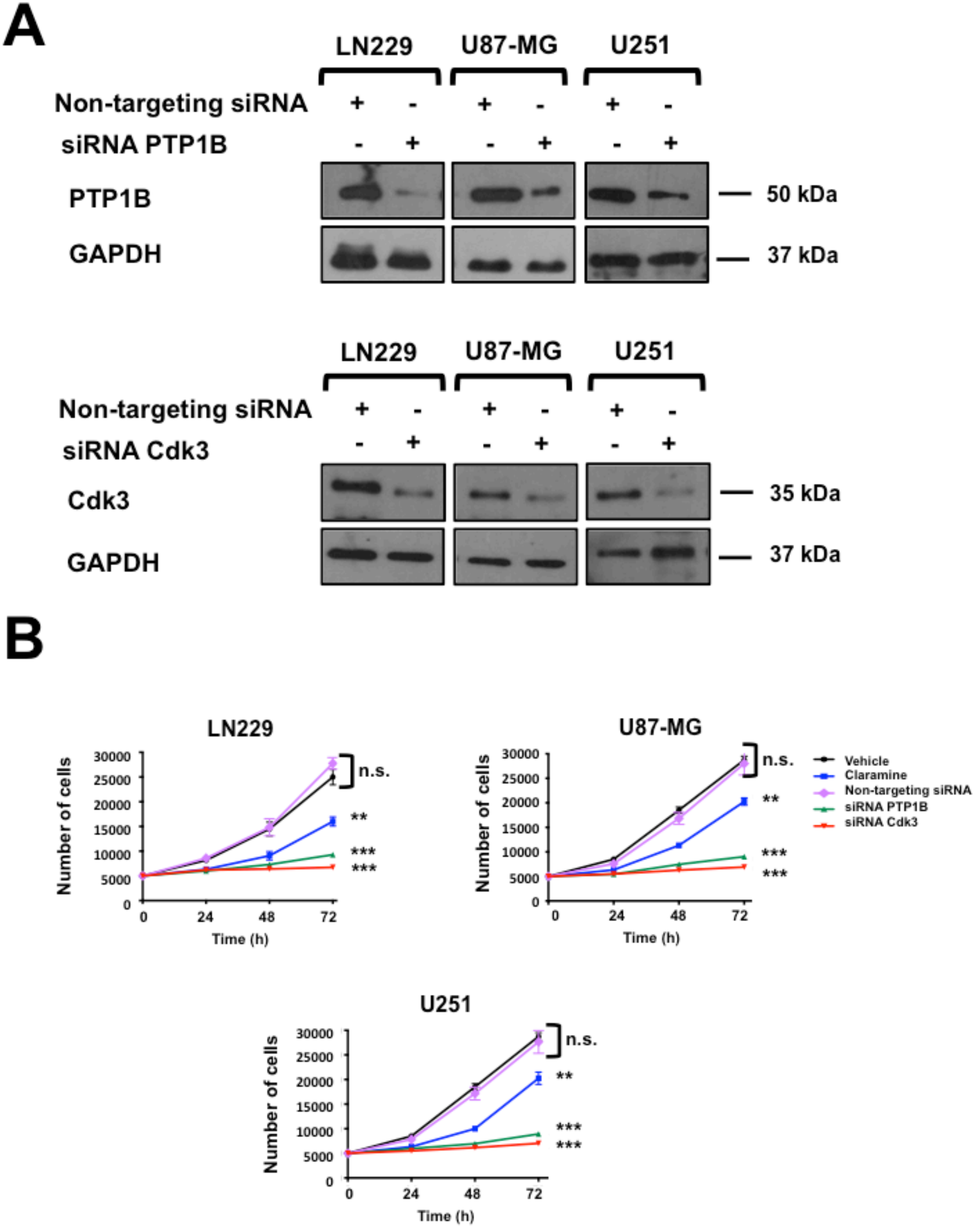


#### Supplementary Figure 2
